# Identification of a novel lineage bat SARS-related coronaviruses that use bat ACE2 receptor

**DOI:** 10.1101/2021.05.21.445091

**Authors:** Hua Guo, Ben Hu, Hao-rui Si, Yan Zhu, Wei Zhang, Bei Li, Ang Li, Rong Geng, Hao-Feng Lin, Xing-Lou Yang, Peng Zhou, Zheng-Li Shi

**Author notes:** These authors contributed equally (Hua Guo, Ben Hu, Hao-rui Si). **Author contributions** Z-L.S and P.Z. conceived the study. X-L.Y organized sampling. B.H., H-R.S, B.L. and Y.Z. performed viral genome sequencing and bioinformatics analysis. H.G.,W.Z., A.L. and R.G. performed protein expression and RBD-ACE2 binding assays. H.G. and H-F.L. performed pseudovirus work. H.G, H.B, P.Z and Z-L.S wrote the paper with input from all authors.

## Abstract

Severe respiratory disease coronavirus-2 (SARS-CoV-2) causes the most devastating disease, COVID-19, of the recent century. One of the unsolved scientific questions around SARS-CoV-2 is the animal origin of this virus. Bats and pangolins are recognized as the most probable reservoir hosts that harbor the highly similar SARS-CoV-2 related viruses (SARSr-CoV-2). Here, we report the identification of a novel lineage of SARSr-CoVs, including RaTG15 and seven other viruses, from bats at the same location where we found RaTG13 in 2015. Although RaTG15 and the related viruses share 97.2% amino acid sequence identities to SARS-CoV-2 in the conserved ORF1b region, but only show less than 77.6% to all known SARSr-CoVs in genome level, thus forms a distinct lineage in the *Sarbecovirus* phylogenetic tree. We then found that RaTG15 receptor binding domain (RBD) can bind to and use *Rhinolophus affinis* bat ACE2 (RaACE2) but not human ACE2 as entry receptor, although which contains a short deletion and has different key residues responsible for ACE2 binding. In addition, we show that none of the known viruses in bat SARSr-CoV-2 lineage or the novel lineage discovered so far use human ACE2 efficiently compared to SARSr-CoV-2 from pangolin or some of the SARSr-CoV-1 lineage viruses. Collectively, we suggest more systematic and longitudinal work in bats to prevent future spillover events caused by SARSr-CoVs or to better understand the origin of SARS-CoV-2.

## Introduction

SARS-CoV-2, a novel coronavirus that causes COVID-19 which was first identified in late 2019 [1], took just a few months to sweep the globe. As the largest pandemic in the past century in human history, it not only results in serious impact on human health but also leads to stagnation in economics, travel, education and many other societal functions globally.

The natural origin of SARS-CoV-2 is one of the unanswered scientific questions about the COVID-19 pandemic. It is generally believed that SARS-CoV-2 is transmitted from an animal reservoir host to human society through an or multiple intermediate hosts [2]. The discovery of SARS-CoV-2 related viruses (SARSr-CoV-2), RaTG13 and Pangolin-CoV from horseshoe bats and pangolin respectively, shed light on the importance of these two groups as animal reservoirs of SARSr-CoV-2 viruses [1,3,4]. However, among the six critical residues of the receptor-binding domain (RBD) in spike to interact with human ACE2 receptor, RaTG13 only shares one with SARS-CoV-2 [5]. The RBD of RaTG13 has a lower binding affinity and usage efficiency with human ACE2 though sharing 96% genome sequence identity to SARS-CoV-2 [6–8]. One of the viruses derived from Malayan pangolin (*Manis javanica*), Pangolin-CoV-GD, possesses six identical critical residues of RBD with SARS-CoV-2 and displays a similar binding affinity to human ACE2 compared with SARS-CoV-2, although it shares lower sequence identity to SARS-CoV-2 in genome compared with RaTG13 [4,7,8]. Another SARSr-CoV-2 detected from bat (*Rhinolophus malayanus*), RmYN02, contains a similar insertion at the S1/S2 cleavage site in the spike of SARS-CoV-2, but it has some deletions in the RBD and fails to bind with human ACE2 [9]. Besides, more SARSr-CoV-2 viral genome sequences from bats have been reported from Eastern China, Japan, and Southeast Asian countries subsequently [10–13]. However, the progenitor virus that shares >99% identical to SARS-CoV-2 was still undetermined.

Bats also carry SARSr-CoV with all the genetic building blocks of SARS-CoV-1, which jumped to human in 2002 [14]. Therefore, investigation of bat SARSr-CoVs is not only important for tracing the origin and immediate progenitor viruses of SARS-CoV-2, but also critical for public health measures to prevent future outbreaks caused by this species of viruses. Here, we report the genome characterization and viral receptor analysis of a novel lineage of SARSr-CoVs in Tongguan town, Mojiang county, Yunnan province in China in 2015, the same location where we found bat RaTG13 in 2013 [1].

## Methods

### Bat Sampling and Coronavirus Detection

Sampling of bat was conducted in Mojiang county, Yunnan province at May 2015. Bats were released after anal swabs sampling. Samples were aliquoted and stored at - 80 °C until use. RNA was extracted using the High Pure Viral RNA Kit (Roche, Basel, Switzerland). Partial RdRp was amplified using the SuperScript III OneStep RT-PCR and Platinum Taq Enzyme kit (Invitrogen, Carlsbad, CA, USA) by family-specific degenerate seminested PCR. The PCR products were gel purified and sequenced with an ABI Prism 3700 DNA analyzer (Applied Biosystems, Foster City, CA). The sequences were blasted against the GenBank database.

### Genome Sequencing

For SARSr-CoV positive RNA extractions, next-generation sequencing (NGS) was performed using BGI MGISEQ 2000. NGS reads were first processed by Cutadapt (v.1.18) to eliminate the possible contamination. Then the clean reads were assembled into genomes using Geneious (v11.0.3) and MEGAHIT (v1.2.9). PCR and Sanger sequencing were used to fill the genome gaps. To amplify the terminal ends, a SMARTer RACE 5’/3’kit (Takara) was used. Bat species identification was based on the partial sequence of cytochrome c oxidase subunit I (COI) gene.

### Phylogenetic Analysis

Routine sequence management and analysis were carried out using DNAStar. Sequence alignments were created by ClustalW implemented in MEGA6 with default parameters. Maximum Likelihood and Neighbour-joining phylogenetic trees were generated using the Jukes-Cantor model with 1000 bootstrap replicates in the MEGA6 software package. Similarity plot analysis of the full-length genome sequences was conducted by Simplot 3.5.1. The genome ID used in the analysis are MN996528 for SARS-CoV-2, AY278488 for SARS-CoV-1, MN996532 for the bat SARSr-CoV RaTG13, MG772933 for ZC45, MW251308 for RacCS203, LC556375 for Rc-o319, KF367457 for WIV1, DQ022305 for HKU3-1, MT121216 for pangolin-CoV-GD strain, MT072864.1 for pangolin-CoV-GX strain, EPI_ISL_412977 for bat SARSr-CoV RmYN02, EPI_ISL_852604 for RshSTT182. The National Genomics Data Center of China ID for the eight novel lineage SARSr-CoVs are: GWHBAUM01000000-GWHBAUT01000000.

### Expression Constructs, Protein Expression, and Purification

Codon-optimized RBD genes from the following viruses were used (see above genome accession number): SARS-CoV-2 (spike aa 330-583), SARS-CoV-1 (spike aa 317-569), RaTG13 (spike aa 330-583), pangolin-CoV-GD (spike aa326-579), pangolin-CoV-GD (spike aa 330-583), RaTG15 (spike aa 317-566). They were synthesized (Sangon Biotech, Shanghai, China) and placed into the expression vector with an N-terminal signal peptide and an S-tag as described previously [15]. The ectodomains of human ACE2 (aa 19-615, accession number: AB046569) and *R.affinis* ACE2 (aa 19-615, accession number: MT394204) were amplified and cloned into the same expression vector as above.

The RBD and ACE2 proteins used for the BLI binding assay were produced in HEK 293T/17 cells. Cells were transiently transfected with expression plasmids using Lipofectamine 3000 (Life Technologies), washed twice with D-Hanks solution 6 h post-transfection, and followed with culturing in fresh 293T FreeStyle expression medium (Life Technologies) at 37°C in a humidified 5% CO2 incubator. The supernatant were harvested 48 h post-transfection and centrifuged at 4000 × *g* for 10 min at 4°C. Clarified supernatant were purified by S-tag agarose beads and eluted with 3 M MgCl_2_. The purified protein was finally buffered with PBS and quantified using Qubit 2 Fluorometer (Thermo Fisher Scientific), and stored at −80 °C until use.

### Bio-layer Interferometry Binding Assays

Binding assays between RBDs and ACE2 proteins were performed using the Octet RED system (ForteBio, Menlo Park, CA, USA) in 96-well microplates at 30°C with shaking at 1000 rpm as described previously [15]. Briefly, the RBD was biotinylated using EZ-Link NHS-LC-LC-Biotin (Thermo Fisher Scientific, Waltham, MA, USA).

The Streptavidin Biosensors were activated for 200s prior to coupling with 50 μg/mL biotinylated RBD proteins for 600s. A baseline were collected in the kinetic buffer (1 M NaCl, 0.1% BSA, 0.02% Tween-20; pH 6.5) for 200s before immersing the sensors in a 1:2 serial diluted ACE2 proteins for 900s and then dissociation in the same kinetic buffer for another 900s. Data analysis from the ForteBio Octet RED instrument includes reference subtraction. Inter-step correction and Y-alignment were used to minimize tip-dependent variability. Curve fitting were performed in a 1:1 model using the Data Analysis Software v7.1 (ForteBio, Menlo Park, CA, USA). The mean Kon, Koff values were determined with a global fit applied to all data. The coefficient of determination (R^2) for these interactions was close to 1.0.

### Pseudovirus Entry Assays

Pseudotyped VSV-^Δ^G particles were generated as previously described with minor adjustments [16]. Briefly, HEK 293T/17 cells were seeded at 6-well-plate and transfected with plasmids contain codon-optimized SARSr-CoV-2 spike at a 70% confluency using Lipofectamine 3000. At 6 h post-transfection, the medium was replaced with fresh DMEM+10%FBS medium. At 24 h after transfection, cells were incubated with VSV-G-pseudotyped VSVG.Fluc at 37°C for 1 h. Cells were subsequently washed five times and supplied with fresh DMEM + 10% FBS medium + anti-VSV-G antibody (Kerafast). Cell-free supernatants were harvested at 24 h after transduction, then centrifuged at 4000 × *g* for 10 min at 4°C. The virus particles were used for infection directly.

The 48-well-plate was treated with Poly-L-lysine solution (Sigma) before seeded HEK293T/17 cells. Cells were transient transfected with equal amounts of human ACE2, *R.affinis* ACE2 or empty vector plasmids at 70% confluency. At 24 h posttransfection, the cells were incubated with same amounts of S-pseudotyped virions for 1 h at 37°C, then washed twice with PBS solution, and supplemented with DMEM containing 10% FBS. Luciferase activity was determined using a GloMax luminometer (Promega Biotech Co. Ltd., Beijing, China) 48 h after infection. Infection experiments were performed independently in triplicate with three technical replications each time.

### Quantification of Pseudotyped Virus Particles using RT-PCR

Viral RNA of all VSV-spike pseudovirus particles were extracted from 200ul supernatant using the High Pure Viral RNA Kit (Roche, Cat. No. 11858882001) following the supplier’s manual. Quantification of pseudovirus by real-time PCR was performed using HiScript^®^ II One Step qRT-PCR SYBR Green Kit (Vazyme, Cat. No. Q221-01). The gene of VSV P protein were amplified and synthesized *in vitro* using mMESSAGE mMACHINE^®^ Kit (Life technologies, Cat. No. AM1344) to serve as a standard. Viral copy numbers were calculated according to the standard curve. Primers using for transcription *in vitro* were: VSV (P protein)-F1: GTTCGTGAGTATCTCAAGTCCT, VSV (P protein)-R2-T7: TAATACGACTCACTATA*GGGAGA*GCCTTGATTGTCTTCAATTTCTGG, primers using for real-time PCR were described as previously [17].

## Results

### Identification of a novel lineage of SARSr-CoVs

In tracing the origin of SARS-CoV-2 from bats, we identified RaTG13, which shares 96.2% genome identity to SARS-CoV-2 and is so far the closest genome [1]. Following the investigation, we identified eight SARSr-CoV sequences that share 93.5% sequence identity to SARS-CoV-2 in the 402-nt partial RdRp gene from bat samples collected at the same place in 2015. Seven samples were from *Rhinolophus stheno*, and the other one was from *Rhinolophus affinis* (Table S1). We thus performed next-generation sequencing (NGS) for further analysis of these CoVs. Whole genome sequences were obtained from all eight individual samples. The eight SARSr-CoV genomes are almost identical, sharing more than 99.7% sequence identity among each other. One strain designated RaTG15 was used as the representative in the subsequent analysis.

In the seven conserved replicase domains used for coronavirus species classification, RaTG15 is 95.3% or 92.5% identical to SARS-CoV-2 and SARS-CoV-1 respectively, suggesting that it remains a member of the *SARSr-CoV* species in the *Sarbecovirus* subgenus within *Betacoronavirus* genus, *Coronaviridae* family. Further, RaTG15 is genetically close to SARS-CoV-2 in open reading frame 1b (ORF1b). In the complete ORF1b region, RaTG15 showed 84.6~89.0% nucleotide identities and 95.6~97.3% amino acid sequence identities to bat SARSr-CoV-2 from wildlife in China and Southeast Asia, which includes bat CoVs RaTG13 and RmYN02 from Yunnan, Rc-o319 from Japan, RshSTT182 from Cambodia, RacCS203 from Thailand, as well as two different strains of pangolin-CoVs (Table S2). It is also conceivable that RaTG15 clusteres with SARSr-CoV-2 in the phylogeny using full-length RdRp gene (Figure S1A).

In contrast, similarity plot analysis reveals that beyond ORF1b, RaTG15 is remarkably distinct from both SARSr-CoV-2 and SARSr-CoV-1 in majority of the genome (Figure 1A). It exhibits less than 80% nucleotide identities in ORF1a, M and N genes and lower than 70% identities in S, ORF3, 6 and 7a/7b to all other SARSr-CoVs (Table S2). Overall, the full genome of the SARSr-CoV RaTG15 show 74.4% sequence identity to SARS-CoV-1 and 77.6% to sequence identity to SARS-CoV-2. Notably, RaTG15 show higher sequence identity to SARS-CoV-1 than to SARS-CoV-2 in the spike, E, M, N and ORF6 proteins. It also has almost equivalent homology to any other known SARSr-CoVs from bat or pangolin CoVs (Table S2). This mosaic profile suggests that this novel lineage viruses may be a results of recombination of different SARSr-CoVs.

**Figure 1.**
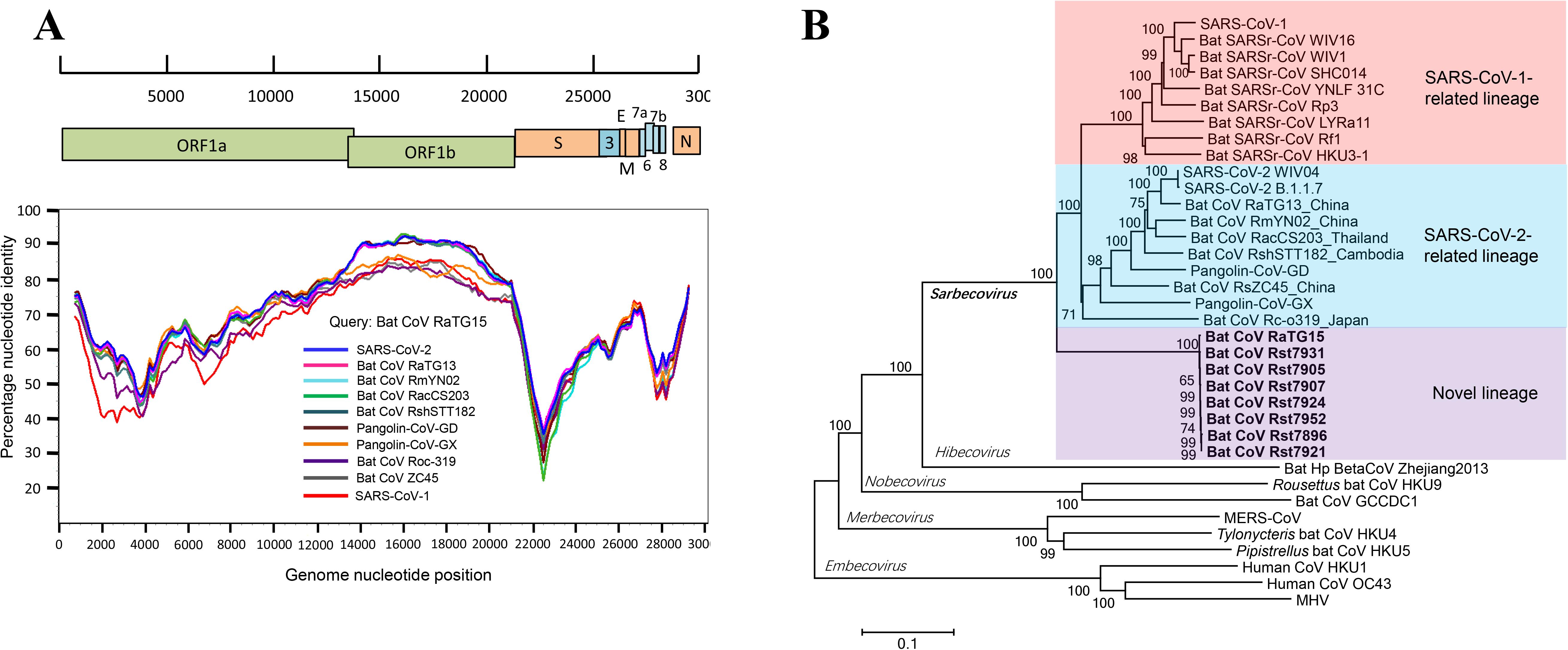
Discovery of a novel lineage of bat SARSr-CoVs. (A) Similarity plot analysis based on the full-length genome sequence of bat SARSr-CoV RaTG15. Full-length genome sequences of SARS-CoV-1, SARS-CoV-2, bat and pangolin CoVs related to SARS-CoV-2 were used as reference sequences. The analysis was performed with the Kimura model, a window size of 1500 base pairs and a step size of 150 base pairs. (B) Phylogenetic tree based on complete genome sequences of betacoronaviruses. The trees were constructed by the Neighbour-joining method using the Jukes-Cantor model with bootstrap values determined by 1000 replicates. Bootstraps > 50% are shown. The scale bars represent 0.1 substitutions per nucleotide position. The novel SARSr-CoVs characterized in this study are shown in bold. Ra, *Rhinolophus affinis*; Rst, *Rhinolophus stheno*; Rsh, *Rhinolophus shameli*; Rs, *Rhinolophus sinicus*; Rac, *Rhinolophus acuminatus*; Rm, *Rhinolophus malayanus*; Rc, *Rhinolophus cornutus*; MHV, murine hepatitis virus.

The result of phylogenetic analysis is in accordance with similarity plot. SARSr-CoVs mainly consists of two sub-lineages, the SARSr-CoV-1 and SARSr-CoV-2 (Figure 1B). The latter one includes SARS-CoV-2 from pangolins and different *Rhinolophus* bats species recently reported in a wide range of areas in Asia. In the full-length genome tree and S gene tree, RaTG15 and the related viruses are distant from both of the two existing sub-lineages, and forms a well-supported novel lineage with the *sarbecoviruses* (Figure 1B and Figure S1B).

### In silico analysis of receptor binding domain (RBDs) of SARSr-CoVs

We further examined the spike protein sequence of RaTG15 in comparison with other SARSr-CoV-2. The receptor-binding domain (RBD) of the RaTG15 spike is highly divergent from other *sarbecoviruses*, with 72.6% amino acid sequence identity to SARS-CoV-2 and 68.6%-73.3% identities to related bat and pangolin CoVs. Unlike RmYN02 and RacCS203, the RaTG15 RBD does not contain the deletion corresponding to aa 473-486 (deletion 2) of the SARS-CoV-2 spike which determines ACE2 usage based on previous reports [18]. However, aligned with SARS-CoV-2 and RaTG13, a short deletion is noted at the position corresponding to aa 444-447 (deletion 1). The location of this deletion is similar to the one in the spike of RshSTT182, a SARS-CoV-2-related CoV identified in *Rhinolophus shameli* from Cambodia. Within the receptor binding motif (RBM), four of the five amino acid residues critical for binding of SARS-CoV-2 to the ACE2 receptor (486, 493, 494 and 501) are varied in RaTG15. Like most bat SARSr-CoVs, the polybasic (furin) cleavage site is absent at the S1-S2 junction of RaTG15 (Figure 2).

**Figure 2.**
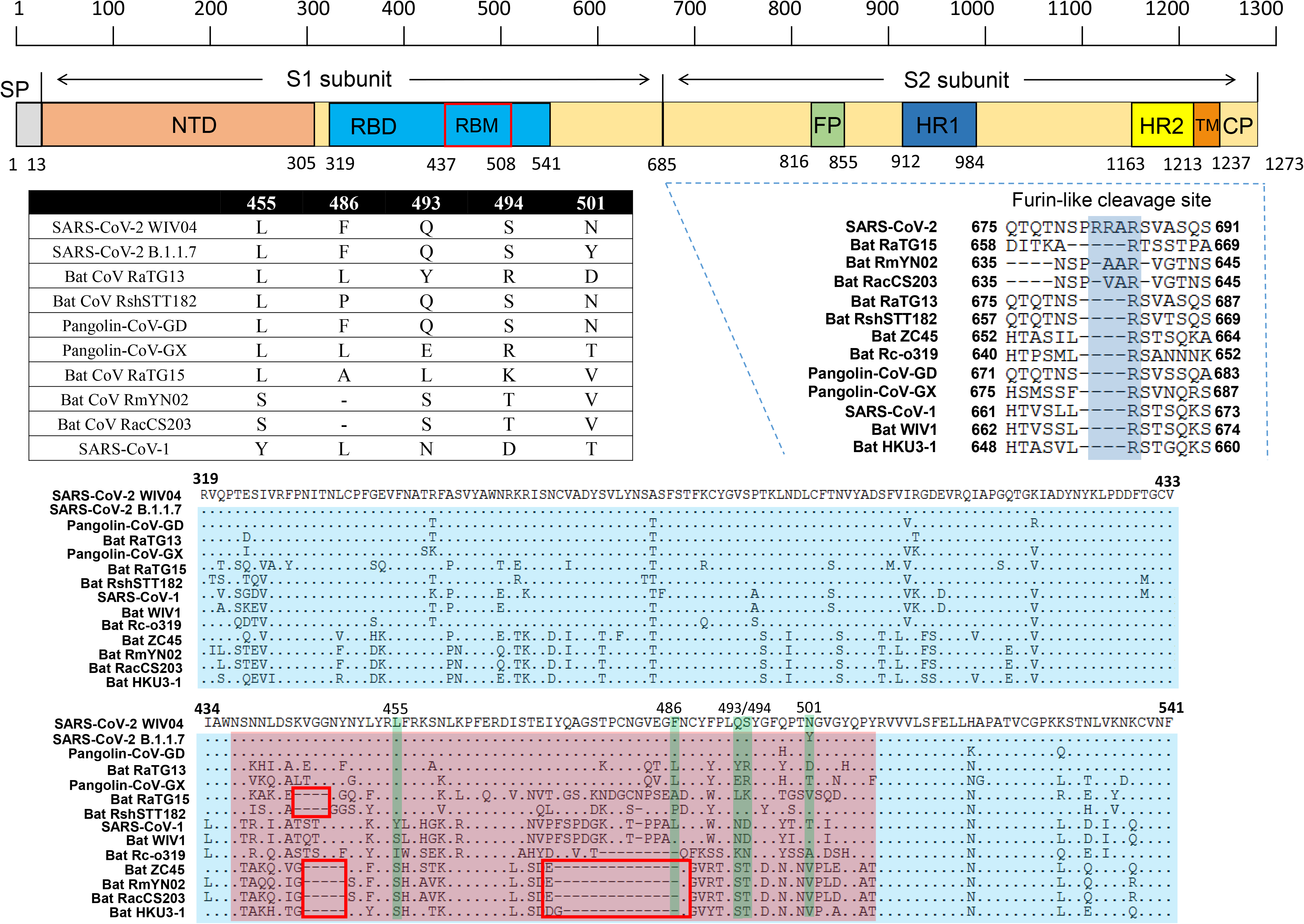
Comparison of receptor-binding domain (RBDs) of SARSr-CoVs. The RBM is shown in pink and the five key residues that contact ACE2 directly are highlighted in green. Comparison of the five critical residues of these SARSr-CoVs are listed in the table. Two deletions in the RBM, aa 444-447 (deletion 1) and aa 473-486 (deletion 2) are indicated by red boxes. GenBank or GISAID entries for each virus can be found in Methods.

### Functional comparison of RBD from three lineages of SARSr-CoVs

The sequence analysis indicated that the RaTG15 virus possibly uses ACE2 as an entry receptor, which was then experimentally confirmed by RBD-ACE2 binding studies using purified recombinant proteins. RBD proteins from SARS-CoV-2, SARS-CoV-1, RaTG13, pangolin-CoV-GD, pangolin-CoV-GX and RaTG15, as well as ectodomains of human and *R.affinis* ACE2 proteins were used (Figure S2A). We found that *R.affinis* derived RaTG13 and RaTG15 RBD proteins either show very weak or have no binding affinity to human ACE2 (HuACE2). In contrast, RBD proteins from the two pangolin SARSr-CoVs displayed much higher binding affinity to HuACE2, only slightly weaker than SARS-CoV-2 RBD but still higher than SARS-CoV-1 (Figure 3A-F and M). Furthermore, the binding affinity to HuACE2 of pangolin-CoV-GX is slightly weaker than pangolin-CoV-GD. Next, we wanted to find out whether bat CoVs RaTG13 and RaTG15 can use *R.affinis* ACE2 more efficiently than huACE2. Detectable binding was observed between RaTG15 RBD and *R.affinis* ACE2 (RaACE2), though the affinity was still weaker than SARS-CoV-2 and pangolin-CoV-GD/GX to RaACE2. RaTG13 RBD showed a very weak binding to RaACE2, same as to HuACE2 (Figure 3G-M and Figure S2B).

**Figure 3.**
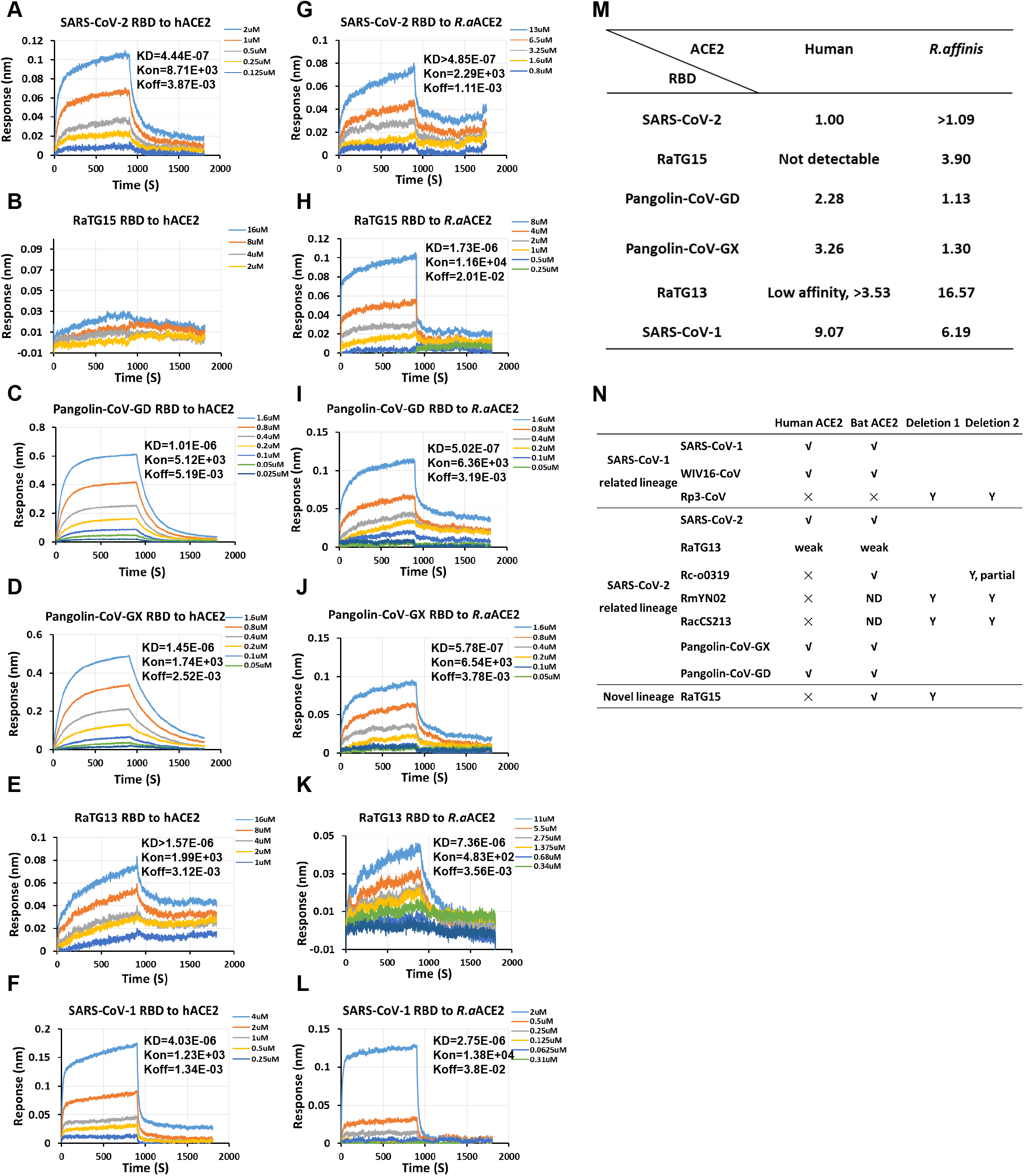
Binding affinity of SARSr-CoV RBDs to ACE2 from human and *R.affinis* bat. (A-F) Binding of different RBD proteins to human ACE2. (G-L) Binding of different RBD proteins to *R.affinis* ACE2. (M) Comparison of dissociation constants (KD) between different RBD to human and *R.affinis* ACE2. Relative binding is analyzed by comparing with SARS-CoV-2 RBD to human ACE2. (N) Summary of the binding efficiency of different RBD to human or bat ACE2. Y, yes; ND, not determined. Evidences for WIV16-CoV, Rc-o0319, RmYN02 and RacCS213 were from previous reports [10,11,21]. The presence of deletion in RBM (related to Figure 2) is indicated. Binding assay of human or *R.affinis* ACE2 to different RBD proteins was measured by Bio-layer interferometry. The parameters of KD value (M), Kon (1/M.s), Koff (1/s) are shown on the upper right side of the picture. Different RBD proteins were immobilized on the sensors and tested for affinity with graded concentrations of human or *R. affinis* ACE2s. The Y-axis shows the real-time binding response. Values reported representing the global fit to all data. The coefficient of determination (R^2) for these interactions was close to 1.0 (Figure S2B).

To exclude the possibility that the ACE2 binding of RBD may not represent the functionality of the full-length S protein, we also constructed a VSV-based pseudovirus using previously published method [16]. We produced a list of SARSr-CoVs pseudoviruses, or MERS-CoV pseudovirus as a negative control. HEK293T/17 cells overexpression HuACE2, RaACE2 or empty vector were infected with VSV-based pseudoviruses, and the infection efficiency were determined 48 h after infection. Consistent with the RBD-ACE2 protein binding assays, HuACE2 mediated entry of all SARSr-CoVs except the RaTG15, whereas the *R.affmis* ACE2 supported all SARSr-CoVs entry. Notably, RaTG13 pseudovirus infection of HuACE2 or RaACE2-expression cells was minimal, if it is positive, compared to other groups. As control, MERS-CoV pseudovirus failed to infect ACE2-expression cells, confirming ACE2-independent infectivity of VSV backbone (Figure S3). Collectively, none of the SARSr-CoV-2 lineage or the novel lineage virus from bats could efficiently bind to HuACE2 [10,11], and it appears that whether there is deletion at RBD region greatly affecting the binding capacity (Figure 3N). These results suggest that without further adaptation, there is a limited zoonotic potential for bat-derived RaTG13, RaTG15 and perhaps other SARSr-CoV-2 lineage or the novel lineage viruses. In contrast, there is a high spillover potential of pangolin-CoV in the context of cell receptor usage.

## Discussion

Overall, we report the discovery of a novel lineage of SARSr-CoVs from bats that are closely related to SARS-CoV-2 in the RdRp region, but genetically distant to any known SARSr-CoVs at genome level. Although several SARS-CoV-2 related coronaviruses have been detected from wildlife, none of them shared >99% genetically identical to SARS-CoV-2 at the genome level. Recombination events happen commonly in coronaviruses and can be referred to as potential origin of the progenitor of SARS-CoV-1, as SARSr-CoVs discoved in a bat colony carried all the genomic fragments of SARS-CoV-1 [14,19]. The high sequence similarity to SARS-CoV-2 in some genomic regions detected from different wildlife species implies the recombination may happen during the virus evolution in cross-species or inter-species transmission. The new lineage virus we reported in this study showed weak binding affinity to bat but not human ACE2 though possess one deletion in the RBD of the spike which is different from the previously reported SARSr-CoVs in bat (Figure 2). These results suggested the SARSr-CoVs we discovered from bat now may be just the tip of the iceberg. These viruses may have experienced selection or recombination events in the animal hosts and render viral adaption to a new host then spread to the new species before they jumped into human society. So surveillance to this new lineage virus should be conducted to prevent future outbreaks, as viruses from the other two lineages of SARSr-CoV caused SARS and COVID-19, respectively [1,20]. Furthermore, none of the bat SARSr-CoV-2 lineage or the novel lineage viruses discovered so far could be isolated, or be capable of efficiently using human ACE2, thus pose little spillover potential to human without future adaptation [21]. In comparison, the ACE2 usage virus in bat SARSr-CoV-1 related lineage appears to be more dangerous in the context of cross-species transmission, which has been demonstrated in animal studies [22,23].

The closest bat CoV to SARS-CoV-2 at this stage, RaTG13 only showed very weak binding affinity to HuACE2. Albeit there is a speculation claiming the possible leaking of RaTG13 from lab that caused SARS-CoV-2, the experiment evidence cannot support it. In contrast, the pangolin-CoV shows strong binding capacity to human or bat ACE2, posing high cross-species potential to human or other species. In the context of SARS-CoV-2 animal origin, there could either be a bat SARSr-CoV closer than RaTG13 that is capable of using HuACE2, or be a pangolin-CoV that obtained higher genome similarity other than spike gene. In future, more systematic and longitudinal sampling of bats, pangolins or other possible intermediate animals is required to better understand the origin of SARS-CoV-2.

## Acknowledgements

We thank Yun-Zhi Zhang, Ji-Hua Zhou from Yunnan CDC for helping with bat sampling. We also would like to thank Dr. Ding Gao in the WIV Core Facility and Technical Support for his help in Octet RED technology. The work was jointly supported by the Strategic Priority Research Program of the Chinese Academy of Sciences (XDB29010101, to Z-L.S) and China National Science Foundation for Excellent Scholars (81290341 to Z-LS, 81822028 to P.Z.).

---

**Figure S1. Phylogenetic tree base on the complete S gene sequences (A) or complete RdRp gene sequences (B) of betacoronaviruses.** The trees were constructed by the Maximum-likelihood method using the Jukes-Cantor model with bootstrap values determined by 1000 replicates. Bootstraps > 50% are shown. The scale bars represent 0.1 and 0.05 substitutions per nucleotide position, respectively. The novel SARSr-CoVs characterized in this study are shown in bold. Ra, *Rhinolophus affinis*; Rst, *Rhinolophus stheno*; Rsh, *Rhinolophus shameli*; Rs, *Rhinolophus sinicus*; Rac, *Rhinolophus acuminatus*; Rm, *Rhinolophus malayanus*; Rc, *Rhinolophus cornutus*; MHV, murine hepatitis virus.

**Figure S2. Binding affinity of SARSr-CoVs RBD proteins to ACE2 from human and *R.affinis*. (A)** The purity of different CoV-RBD and ACE2 proteins used for binding assay were analyzed by SDS-PAGE. **(B)** Binding assay of human or *R.affinis* ACE2 to different RBD proteins measured by Bio-layer interferometry, Related to **Figure 3.** The Y-axis shows the real-time binding response. Values reported representing the global fit to all data. The coefficient of determination (R^2) for these interactions was shown on the upper right.

**Figure S3. Infectivity analysis of SARSr-CoV spike VSV-pseudoviruses in human and *R.affinis* ACE2 expression cells.** HEK293T/17 cells expression human/*R.affinis* ACE2 were infected with SARS-CoV-2 and SARSr-CoV spike-pseudotyped viruses. The infected cell lysis was analyzed by measuring luciferase activities. All results were performed in triplicate from three independent experiments. Error bars indicate mean ± SEM. Statistical significance was tested by one-way ANOVA with Dunnett posttest. (A) SARS-CoV-2, (B) RaTG15, (C) pangolin-CoV-GD, (D) pangolin-CoV-GD, (E) RaTG13, (F) SARS-CoV-1, (G) MERS-CoV. (H) Genome copies of VSV-CoV-S pseudotyped particles. Viral copy numbers were calculated according to the standard curve of VSV P protein gene. A representative result is shown. (I) ACE2 expression was detected using mouse anti-Stag monoclonal antibody followed by HRP-labelled goat anti-mouse IgG antibody. β-actin was detected with mouse anti-β-action monoclonal antibody by HRP-labelled goat anti-mouse IgG antibody.

