## Supplementary figures and images for "Identification of a novel lineage bat SARS-related coronaviruses that use bat ACE2 receptor"

### Fig S1

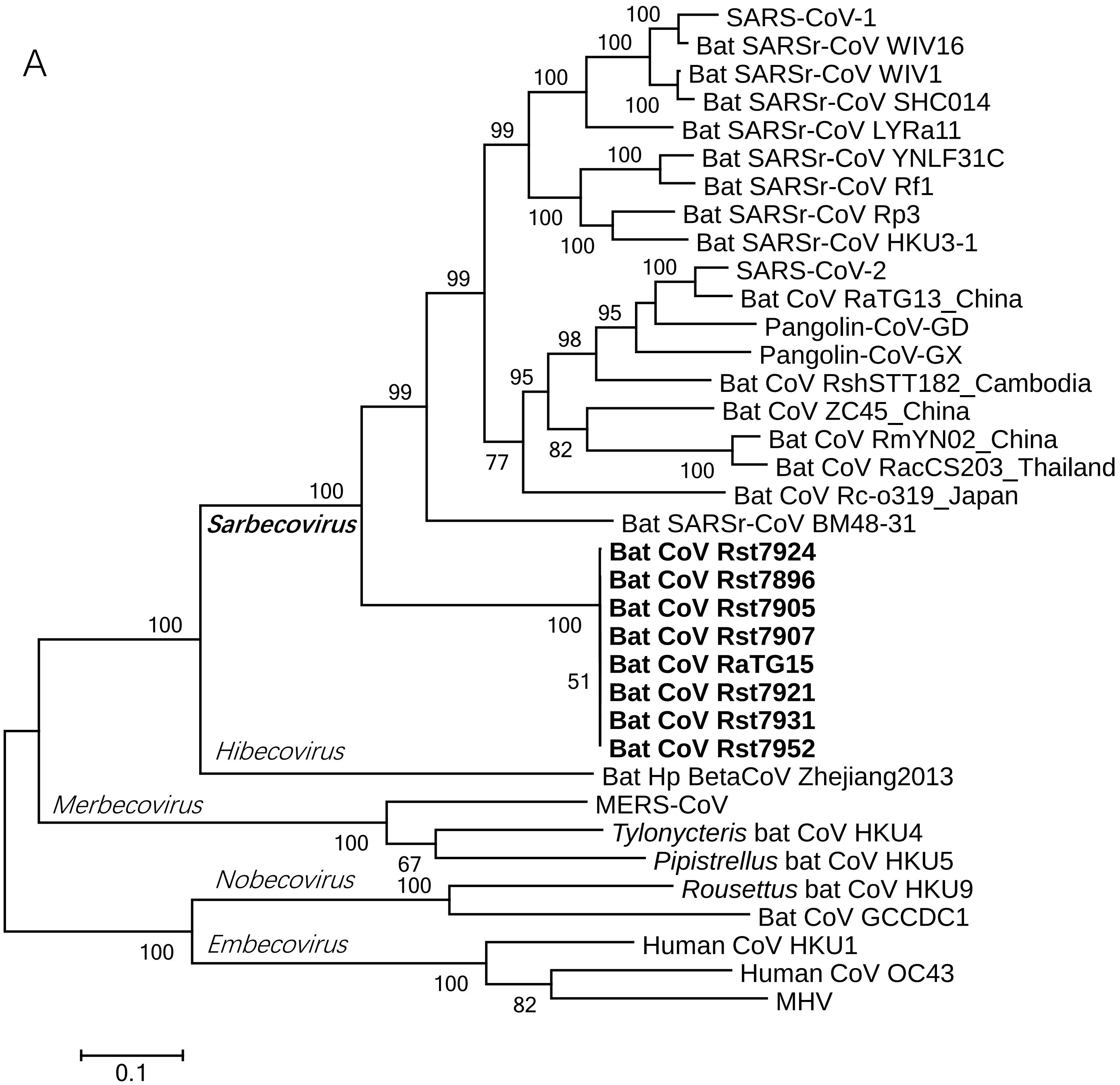

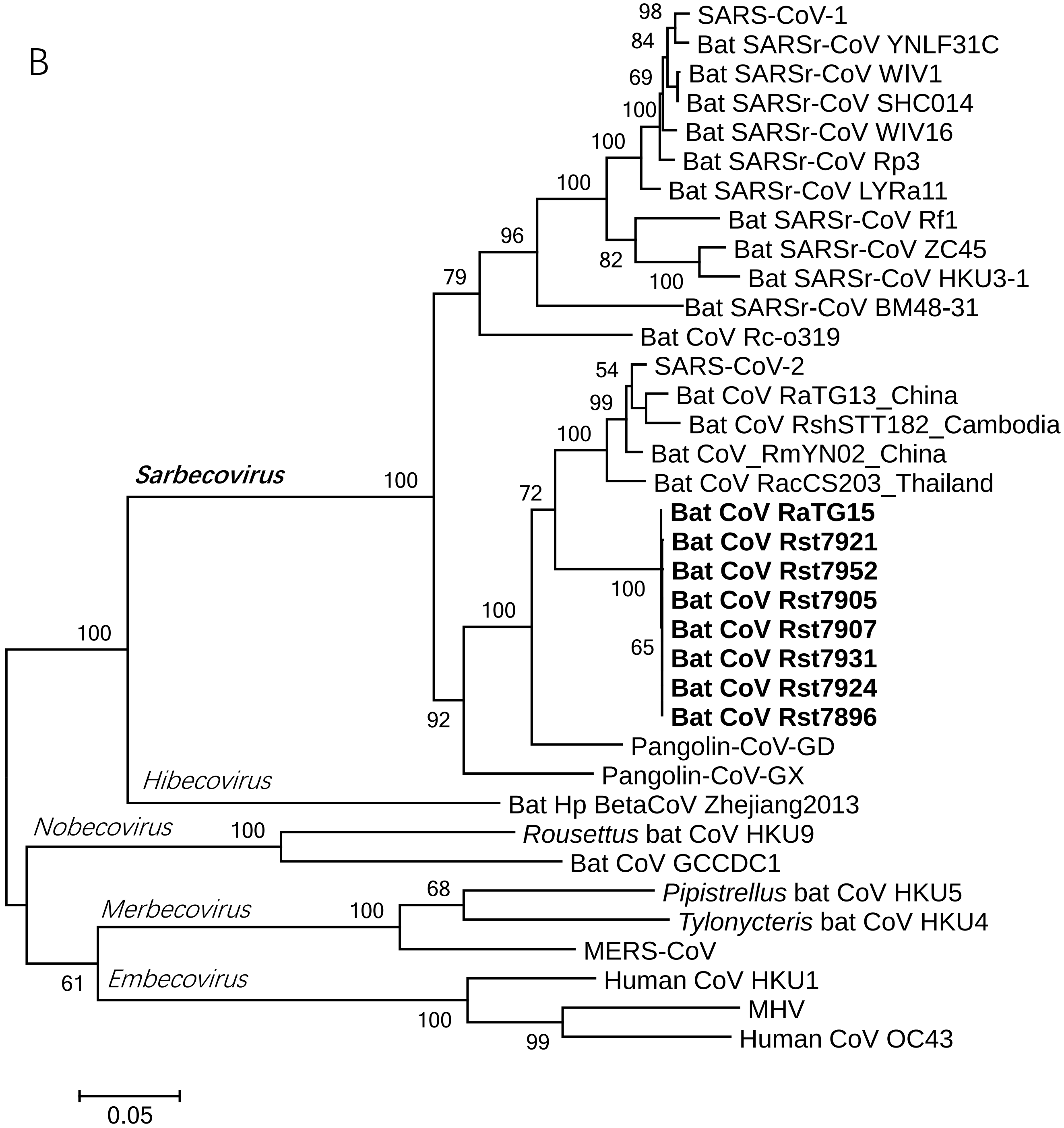

### Fig S2

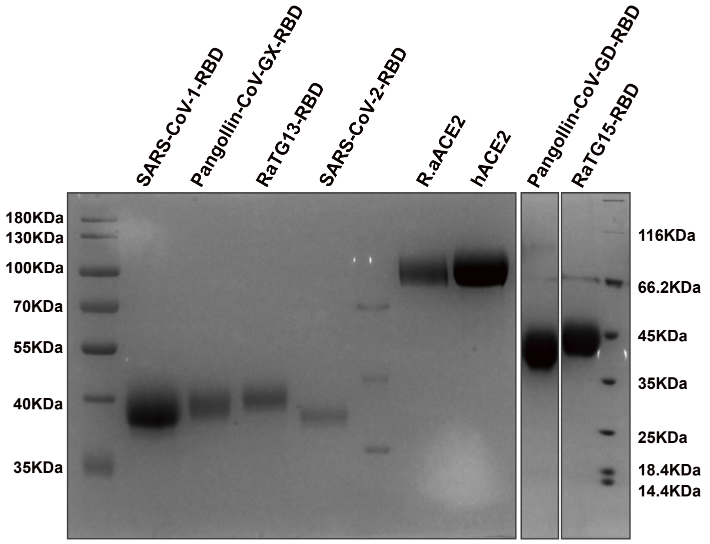

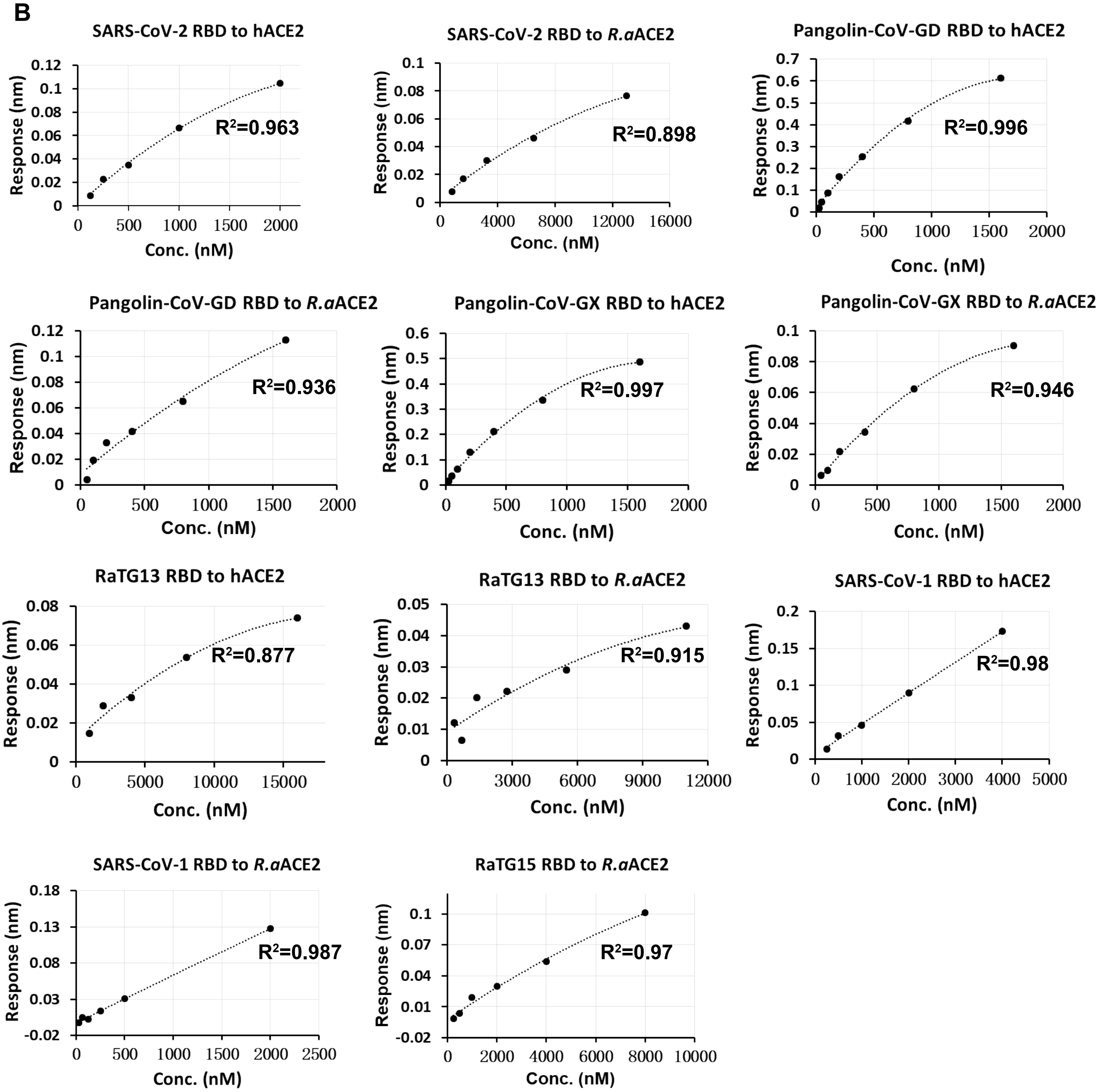

### Fig S3

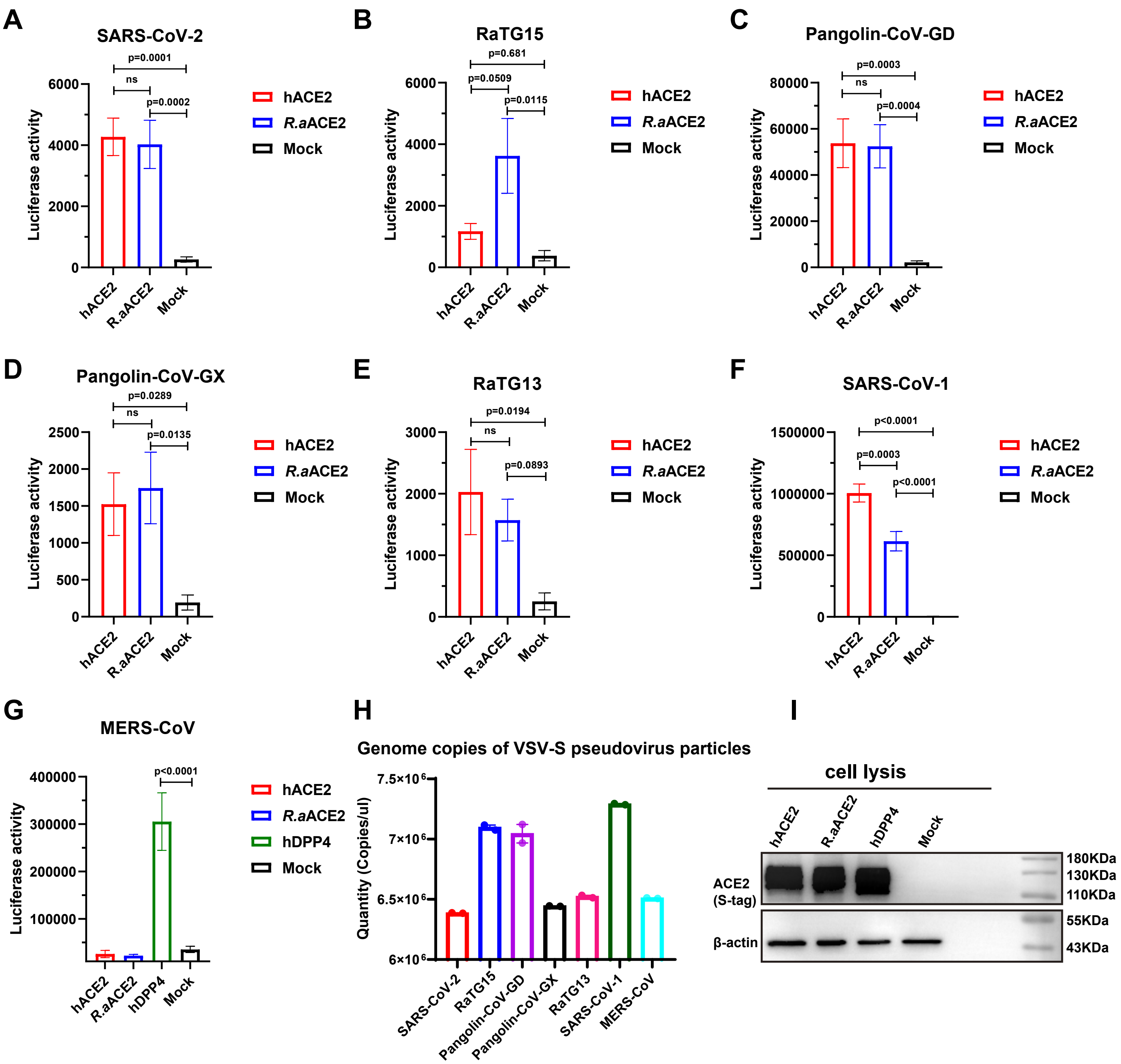
