## Supplementary material for "Identification of a novel lineage bat SARS-related coronaviruses that use bat ACE2 receptor": Table S1-S2

**Supplemental Information**

**Table S1**. Sampling information of Bat CoV RaTG15 and related CoVs

| **Sample ID** | **Sampling date** | **Sampling location** | **GPS information** | **Host bat species** |
| --- | --- | --- | --- | --- |
| 7896 | 20150529 | Tongguan town, Mojiang county, Yunnan province, China | N 23°3'27073"，E 101°37'16074" | Rhinolophus stheno |
| 7905 | 20150529 | Tongguan town, Mojiang county, Yunnan province, China | N 23°3'27073"，E 101°37'16074" | Rhinolophus stheno |
| 7907 | 20150529 | Tongguan town, Mojiang county, Yunnan province, China | N 23°3'27073"，E 101°37'16074" | Rhinolophus stheno |
| RaTG15 (7909) | 20150529 | Tongguan town, Mojiang county, Yunnan province, China | N 23°3'27073"，E 101°37'16074" | Rhinolophus affinis |
| 7921 | 20150529 | Tongguan town, Mojiang county, Yunnan province, China | N 23°3'27073"，E 101°37'16074" | Rhinolophus stheno |
| 7924 | 20150529 | Tongguan town, Mojiang county, Yunnan province, China | N 23°3'27073"，E 101°37'16074" | Rhinolophus stheno |
| 7931 | 20150529 | Tongguan town, Mojiang county, Yunnan province, China | N 23°3'27073"，E 101°37'16074" | Rhinolophus stheno |
| 7952 | 20150529 | Tongguan town, Mojiang county, Yunnan province, China | N 23°3'27073"，E 101°37'16074" | Rhinolophus stheno |

**Table S2**. Genomic comparison of Bat CoV RaTG15 with SARS-CoV-2, SARS-CoV-1 and their related CoVs

| Sequence identities with SARS-CoV-2, SARS-CoV-1 and related bat and Pangolin CoVs (nt/aa %) | | | | | | | | | | | | | |
| --- | --- | --- | --- | --- | --- | --- | --- | --- | --- | --- | --- | --- | --- |
|  | Full-length genome | | | ORF1a | ORF1b | S | ORF3 | E | M | ORF6 | ORF7a | ORF7b | N |
| SARS-CoV-2 | | 77.6 | | 74.9/80.5 | 88.9/97.2 | 64.4/68.4 | 69.6/66.1 | 86.0/80.0 | 78.0/90.0 | 69.5/57.9 | 64.2/54.2 | 53.8/37.2 | 78.7/86.6 |
| Bat CoV RaTG13 | 77.5 | | | 74.8/80.6 | 88.7/96.8 | 64.9/68.6 | 69.4/66.8 | 85.5/80.0 | 77.2/89.5 | 69.0/57.9 | 63.4/53.3 | 53.0/34.9 | 78.7/86.8 |
| Bat CoV RshSTT182 | 77.2 | | | 74.4/79.9 | 88.5/97.0 | 64.8/68.6 | 69.5/66.4 | 85.1/80.0 | 77.5/89.5 | 68.4/57.9 | 65.0/53.3 | 53.0/34.9 | 78.9/87.5 |
| Pangolin-CoV-GD | 77.5 | | | 75.0/80.8 | 89.0/97.3 | 64.1/68.6 | 70.2/67.5 | 85.1/80.0 | 79.2/89.1 | 71.3/57.9 | 64.5/54.2 | 47.7/34.9 | 78.9/87.0 |
| Pangolin-CoV-GX | 76.5 | | | 75.1/80.8 | 84.6/95.6 | 64.7/68.2 | 70.6/68.3 | 86.4/80.0 | 79.2/88.6 | 67.8/59.6 | 60.6/52.5 | 47.0/32.6 | 78.4/86.1 |
| Bat CoV RmYN02 | 77.1 | | | 74.6/80.3 | 88.8/97.0 | 63.0/67.1 | 70.5/66.4 | 86.0/80.0 | 77.7/89.5 | 69.0/56.1 | 64.5/53.3 | 53.0/37.2 | 78.9/86.8 |
| Bat CoV RacCS203 | 77.2 | | | 74.6/80.2 | 88.5/96.6 | 63.5/67.3 | 70.5/67.2 | 85.1/80.0 | 77.8/89.1 | 70.7/57.9 | 64.2/51.7 | 53.8/37.2 | 78.7/87.0 |
| Bat CoV Rc-o319 | 74.5 | | | 72.5/77.7 | 82.6/93.9 | 64.1/68.1 | 69.9/67.2 | 85.5/80.0 | 78.3/91.8 | 66.1/57.9 | 62.6/52.9 | 50.8/32.6 | 78.7/86.3 |
| Bat CoV ZC45 | 75.5 | | | 74.2/80.2 | 83.0/94.4 | 64.5/67.2 | 69.1/64.2 | 85.5/80.0 | 78.7/89.5 | 67.8/59.6 | 63.9/53.3 | 53.8/34.9 | 78.9/86.8 |
| SARS-CoV-1 | 74.4 | | | 71.2/76.7 | 83.6/94.4 | 65.2/69.3 | 69.7/64.9 | 85.7/81.6 | 79.9/92.7 | 67.2/60.3 | 62.6/53.7 | 51.1/36.4 | 80.2/88.5 |
| Bat SARSr-CoV WIV1 | | 74.6 | | 71.2/76.7 | 83.5/94.5 | 64.9/69.8 | 70.1/66.4 | 85.7/81.6 | 79.2/91.8 | 66.7/62.1 | 63.7/54.5 | 52.6/36.4 | 79.9/88.3 |
| Bat SARSr-CoV HKU3-1 | | | 74.4 | 71.1/76.8 | 82.7/94.0 | 65.1/68.0 | 70.9/67.2 | 85.3/81.6 | 80.4/93.2 | 68.4/58.6 | 64.2/55.4 | 51.1/34.1 | 79.4/88.0 |
